# Trade-offs among plant reproductive traits determine interactions with floral visitors

**DOI:** 10.1101/2021.12.09.471959

**Authors:** Jose B. Lanuza, Romina Rader, Jamie Stavert, Liam K. Kendall, Manu E. Saunders, Ignasi Bartomeus

## Abstract

Plant life-history strategies are constrained by cost-benefit trade-offs that determine plant form and function. However, despite recent advances in the understanding of trade-offs for vegetative and physiological traits, little is known about plant reproductive economics and how they constrain plant life-history strategies and shape interactions with floral visitors. Here, we investigate plant reproductive trade-offs and how these drive interactions with floral visitors using a dataset of 17 reproductive traits for 1,506 plant species from 28 plant-pollinator studies across 18 countries. We tested whether a plant’s reproductive strategy predicts its interactions with floral visitors and if the different reproductive traits predict the plant’s role within the pollination network. We found that over half of all plant reproductive trait variation was explained by two independent axes that encompassed plant form and function. Specifically, the first axis indicated the presence of a trade-off between flower number and flower size, while the second axis indicated a pollinator dependency trade-off. Plant reproductive trade-offs helped explain partly the presence or absence of interactions with floral visitors, but not differences in visitation rate. However, we did find important differences in the interaction level among floral visitor guilds on the different axes of trait variation. Finally, we found that plant size and floral rewards were the most important traits in the understanding of the plant species network role. Our results highlight the importance of plant reproductive trade-offs in determining plant life-history strategies and plant-pollinator interactions in a global context.

---

Despite the astonishing diversity of floral structures among flowering plants^1,2^ and their importance in shaping plant-pollinator interactions^3,4^, a unified framework that explores plant reproductive trade-offs is currently lacking^5^. In addition, macroecological studies that investigate plant reproductive traits are scarce^6–9^ and consequently, there is poor understanding of how reproductive traits drive interactions with floral visitors at large scales^10–13^. Linking the plant’s position in trait-space with the different pollinator groups could help to improve our understanding of plant-pollinator associations^14^. Further, there is increasing interest in understanding drivers of plant-pollinator interactions using trait-based approaches^3,15^ and trait-matching analyses^16,17^. However, despite the generalist nature of most plant-pollinator interactions^18,19^, reproductive traits have been overlooked beyond highly specialised pollination systems^4^. Overall, it is unclear how specific plant reproductive biology traits shape plant-pollinator interactions^20,21^.

Species can optimise their fitness through various life-history traits, yet trade-offs among those traits constrain the range of potential strategies that a species can use. With the recent availability of large trait databases (e.g., TRY^22^ and COMPADRE^23^), plant ecological strategies are being increasingly examined, and are facilitating the identification of global patterns and constraints in plant form and function^12,24–26^. However, most studies have focused on vegetative traits such as leaf^27^, wood^28^, or root^29^ trade-offs with little or no attention given to reproductive traits^5,30^ which are critical to plant life strategies that shape interactions with pollinators and ultimately determine plant reproductive success. For instance, short lived versus perennial species tend to have low versus high levels of outcrossing, respectively,^9,31^ and outcrossing levels are positively correlated with flower size^32^. In addition, the presence of costly rewards (e.g., pollen or nectar) and showy flowers or floral displays can only be understood through consideration of plant species’ reliance upon animal pollination (pollinator dependence) and its role in attracting pollinators^33,34^. However, it is still unknown to what extent these different reproductive compromises determine plantpollinator interactions.

Several studies have identified links between plant traits and plant-pollinator network properties^35–37^. Moreover, plant traits can define species’ network roles (e.g., specialists vs generalists)^20,38^. For example, plant species that occupy reproductive trait space extremes are more likely to exhibit higher levels of specialisation and be more reliant on the trait-matching with pollinators^39,40^. Morphological matching between plant and floral visitors often determines plant-pollinator interactions, and can thus strongly influence interaction network structure^16,41^. Remarkably, the combination of traits have shown to increase the predictive power of the network interactions^42^. Therefore, considering the different plant reproductive trade-offs which represent the species reproductive strategy within the network^14^ could progress our understanding of plantpollinator interactions. Further, we know little if those patterns generally studied at the community level are representative of wider macroecological scales.

Here, we aim to explore the potential trade-offs among reproductive traits and how these influence plant-pollinator interactions. First, we identify the major axes of reproductive trait variation and trade-offs that determine plant form and function. Second, we investigate how plant species’ position in trait-space influence interactions with floral visitors. Finally, we investigate how both the main axes of trait variation, and individual traits, influence plant species’ roles within networks using a set of complementary interaction network metrics (i.e., interaction strength, normalized degree and specialization).

## RESULTS

### Plant strategies

The phylogenetically informed principal component analysis (pPCA) captured by the first two and three axes 51.8% and 70.97% of trait variation, respectively (Fig. 1 and Supplementary Fig. S5) and had a phylogenetic correlation (*λ*) of 0.76. The first principal component (PC1) represented 26.72% of the trait variation and indicated a trade-off between flower number and flower size. We refer to this axis as the ‘flower number - flower size trade-off’, as already described in previous studies^43,44^. Hence, one end of the spectrum comprised species with high investment in flower number and plant height but small flower size, short style length and low ovule number. The other end of this spectrum comprised species that were short in height and invested in large flowers, long styles, many ovules, but few flowers. The main contributing traits to PC1 were plant height, flower number, ovule number and flower size (loadings > |0.5|; Supplementary Table S3) but style length also contributed moderately to PC1 (loading = -0.33). The second principal component (PC2) represented 25.05% of the trait variation and indicated a trade-off between low and high pollinator dependence. We refer to this axis as the ‘pollinator dependence trade-off’. The main driver of trait variation on PC2 was autonomous selfing (loading = 0.85) but the other traits (except ovule number) also made moderate contributions (loadings from 0.27 to 0.4; Supplementary Table S3). We found that high pollinator dependence was associated with larger and a higher number of flowers, greater plant height and longer styles. In contrast, species with high levels of autonomous selfing tended to have fewer and smaller flowers, had shorter styles and were shorter in height. Further, PC3 explained a considerable amount of trait variability (19.17%) and the main contributors to this axis were style length (loading = -0.66) and the degree of autonomous selfing (loading = -0.51). The remaining traits, apart from ovule number, were moderately correlated to changes on PC3 (loadings from -0.23 to -0.46; Supplementary Table S3). Thus, because style length was correlated with all traits on PC3 and was the main driver of trait variation, we refer to this axis as the ‘style length trade-off’. Further, the pPCA with the subset of species that had nectar and pollen quantity data showed that nectar quantity (microlitres of nectar per flower) was positively associated with flower size, style length and ovule number (PC1, 23.40%); and pollen quantity (pollen grains per flower) was positively correlated with flower number and plant height and negatively associated with autonomous selfing (PC2, 21.67%; Supplementary Fig. S6). This pPCA explained similar variance with the first two principal components (45.07%) and similar associations of traits despite some variability in the loadings (Supplementary Table S4).

**Fig. 1.**
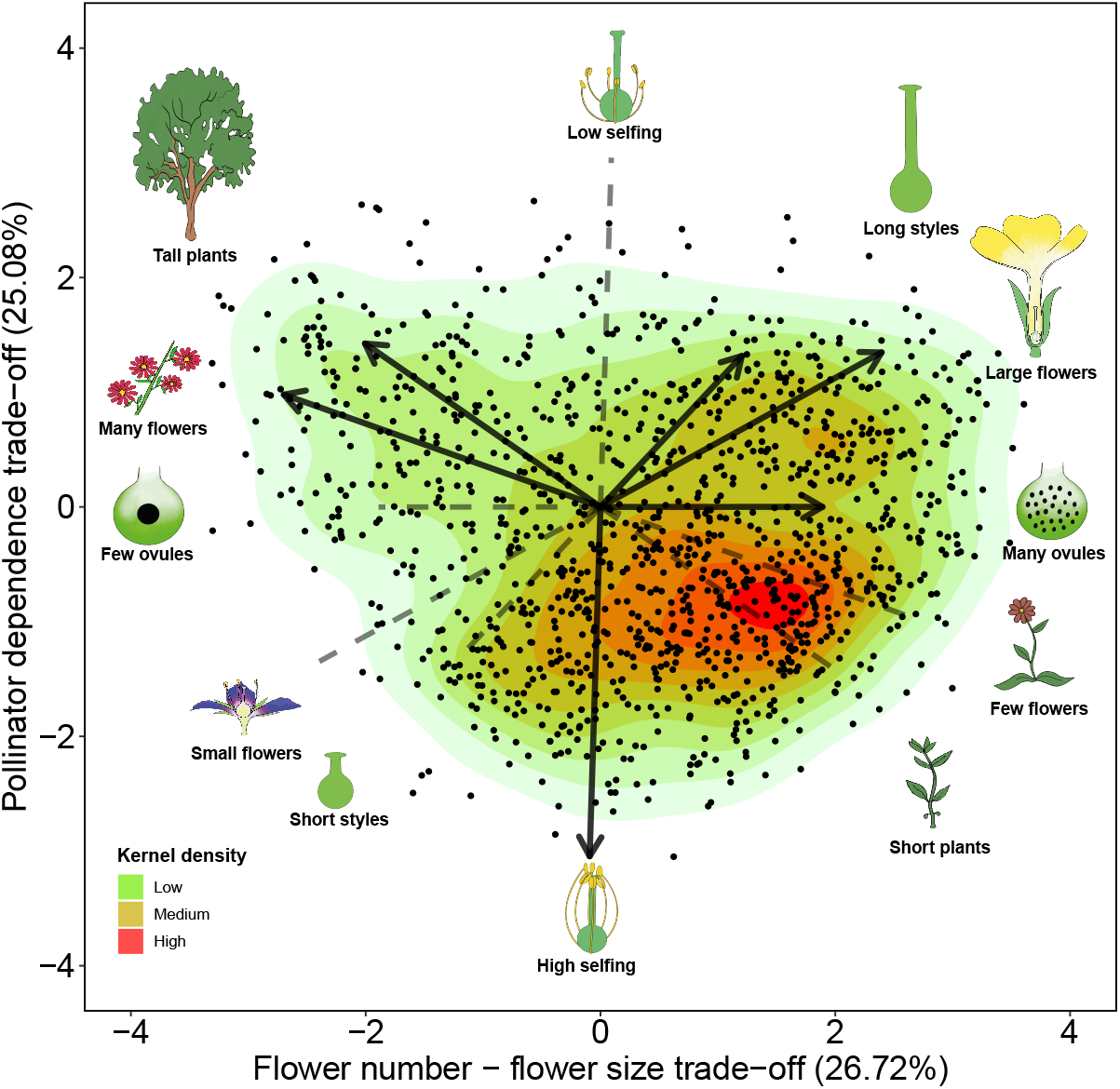
Plant life-history strategies. Phylogenetically informed principal component analysis (pPCA) of 1,236 plant species from 28 plant-pollinator network studies. The solid arrows indicate the direction of the different quantitative traits (flower number, plant height, style length, flower size, ovule number and level of autonomous selfing) across the two main axes of trait variation. The length of the arrows indicate the weight of the variables on each principal component and the dashed lines show the opposed direction of trait variation. The icons at both ends of arrows and dashed lines illustrate the extreme form of the trait continuum.

We found that most categorical traits were statistically associated with the first two axes of trait variation (Fig. 2 and Supplementary Table S2). Flower symmetry, which was only associated with PC2 (Sum of squares = 8.51, F-value = 14.72, *P* < 0.01), and nectar provision, which was independent of PC1 and PC2 (PC1: Sum of squares = 0.37, F-value = 0.29, *P* = 0.59; PC2: Sum of squares = 0.83, F-value = 1.43, *P* = 0.23) showed lack of statistical association. In addition, we found (with a Tukey test) statistical differences between the different levels of categorical traits in the trait space (Supplementary Fig. S7). Regarding self compatibility, we found larger differences on PC2 (i.e., species with unisexual flowers that were self incompatible were statistically differentiated from species with partial or full self compatibility; Supplementary Fig. S7a and Fig. S7b). Life forms differed statistically across both axes of trait variation and followed a gradient of larger life forms (trees and shrubs) with higher pollinator dependence to smaller ones (herbs) with lower pollinator dependence (Supplementary Fig. S7c and Fig. S7d). Consequently, lifespan also followed this gradient but perennial and short lived species only differed statistically on PC2 (Supplementary Fig. S7e and Fig. S7f). Species with unisexual flowers (monoecious and dioecious) were clustered on both extremes of the first two principal components and had the highest pollinator dependence and highest number of flowers (Supplementary Fig. S7g and Fig. S7h). Moreover, we found that the campanulate and capitulum flower shapes were differentiated from tube, papilionaceous, open and brush shapes in the trait space. The former morphologies had larger flowers and greater pollinator dependence, while the latter had higher flower number and greater autonomous selfing (Supplementary Fig. S7i and Fig. S7j). Regarding flower symmetry, zygomorphic flowers were associated with lower levels of pollinator dependence, whereas actinomorphic flowers had higher levels of pollinator dependence (Supplementary Fig. S7k and Fig. S7l).

**Fig. 2.**
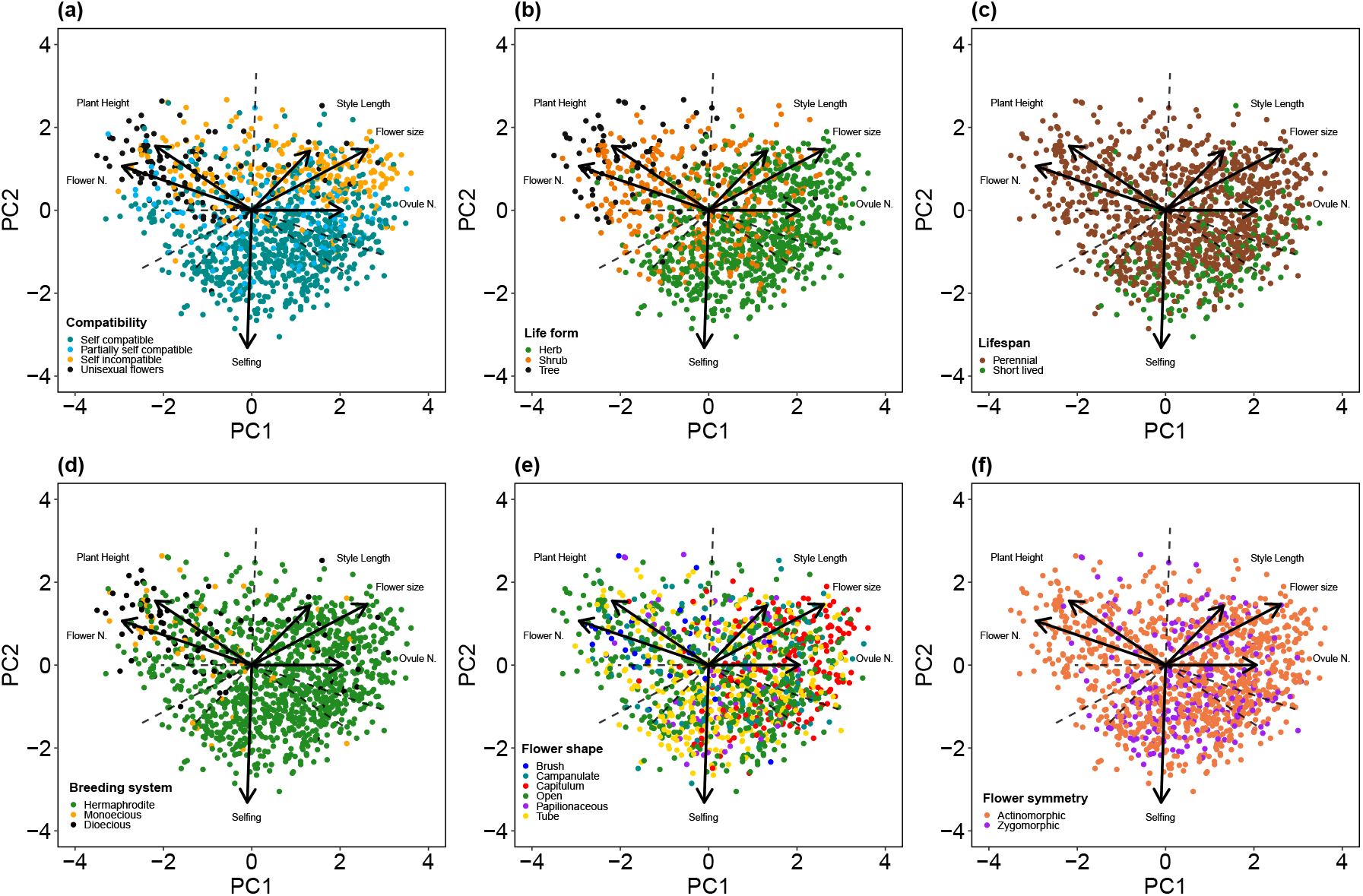
Location of the different qualitative traits on the trait space. The panel is composed by the traits that showed statistical association with the first two axes of trait variation: compatibility system (a), life form (b), lifespan (c), breeding system (d), flower shape (e) and flower symmetry (f).

### Phylogenetic signal of traits

We found a strong phylogenetic signal (*P* < 0.01) in all quantitative traits (Supplementary Table S5). The traits that showed the highest phylogenetic signal were ovule number (*λ* = 1), pollen grains per flower (*λ* = 1) and plant height (*λ* = 0.96), followed by flower length (*λ* = 0.75), flower width (*λ* = 0.73), number of flowers per plant (*λ* = 0.69) and nectar concentration (*λ* = 0.65). The traits that showed a moderate phylogenetic signal were inflorescence width (*λ* = 0.57), style length (*λ* = 0.49) and autonomous selfing (*λ* = 0.34). Finally, microliters of nectar per flower showed the lowest phylogenetic signal of all traits (*λ* = 0.14).

### Visitation patterns

The main axes of trait variation explained partly presence-absence interactions between plant and floral visitors (conditional *R*^2^ = 0.26; marginal *R*^2^ = 0.20) but little of the overall visitation rates (conditional *R*^2^ = 0.31; marginal *R*^2^ = 0.06). However, we found relevant trends across the different floral visitor guilds on both presence-absence and visitation interactions (Fig. 3). On the pollinator dependence trade-off, all floral visitor guilds interacted more frequently with plant species with higher pollinator dependence (PC2; Fig. 3b and Fig. 3e). For presence-absence interactions we found that all Diptera, Coleoptera and non-bee-Hymenoptera guilds interacted more frequently with plants with high flower number and small flowers (flower number - flower size trade-off, PC1; Fig. 3a) but bees and Lepidoptera interacted slightly more frequently with plant species with low flower number but large flowers. For presence-absence interactions on PC3 (style length trade-off; Fig. 3c), we found that bees interacted clearly more with plant species with long styles and high selfing and the rest of the guilds interacted slightly more with plant species with short styles and low selfing. In addition, all guilds other than Syrphids and Lepidoptera (i.e., all Hymenoptera, non-syrphid-Diptera and Coleoptera) showed greater visitation rates on species with small numerous flowers (PC1; Fig. 3d). On the style length trade-off, bees, Lepidoptera and non-bee-Hymenoptera showed greater visitation rates on plant species with larger styles and higher levels of selfing; while syrphids, non-syrphid-Diptera and Coleoptera showed higher visitation rates on species with shorter styles and lower selfing (Fig. 3f).

**Fig. 3.**
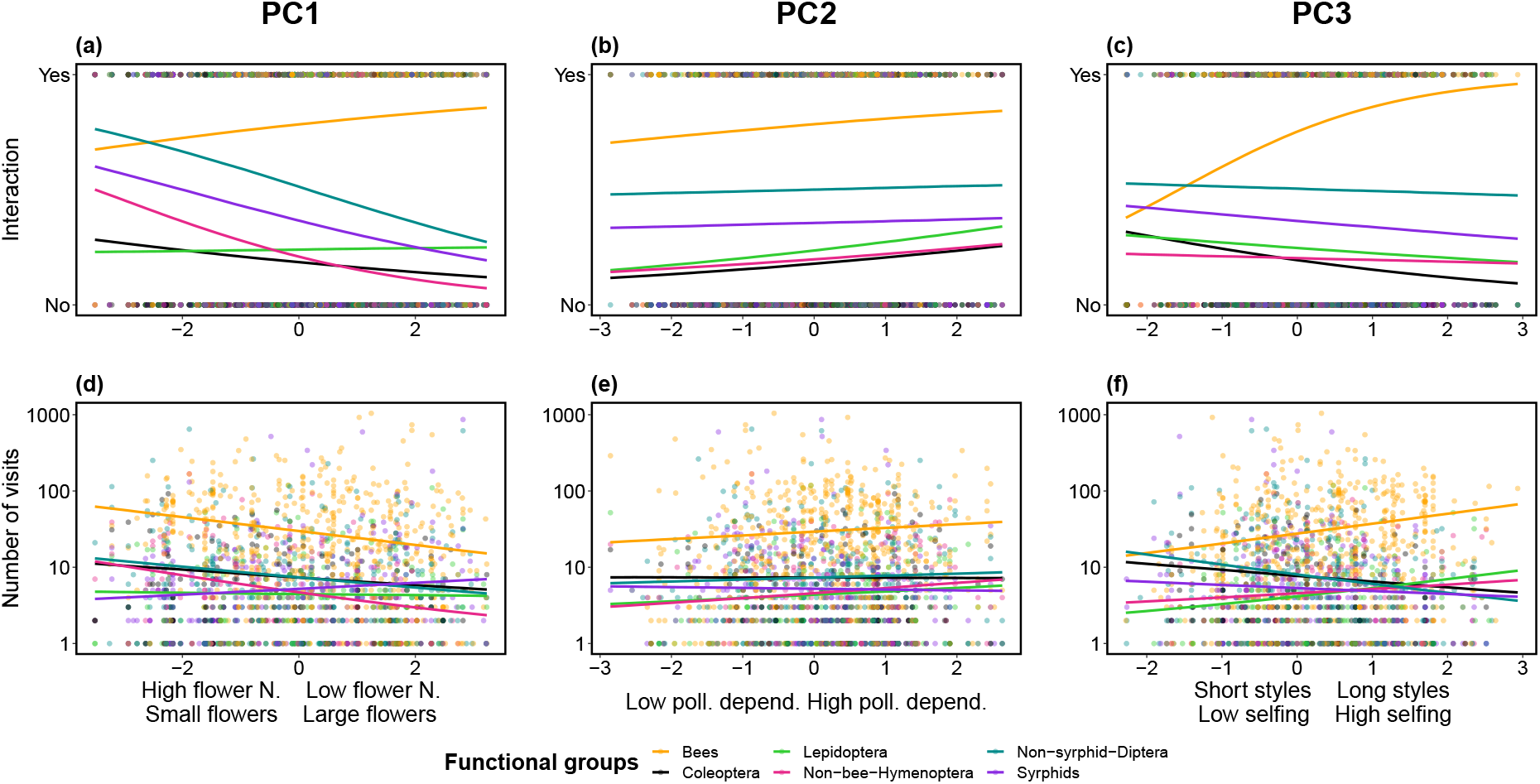
Interaction (yes/no) and visitation rates across the three main axes of trait variation per floral visitor guild. Fitted posterior estimates of the presence/absence of interaction (a, b and c) and number of visits (d, e and f) made by the different floral visitors guilds in relation to PC1, PC2 and PC3. PC1 represents the flower number - flower size trade-off, PC2 represents the pollinator dependence trade-off and PC3, the style length trade-off. For visualization purposes, due to large differences between the visitation rates of bees and the rest of guilds, the number of visits was log-transformed (Y-axis of lower panel).

The additional model for both presence-absence of interactions (marginal *R*^2^ = 0.29; conditional *R*^2^ = 0.19) and visitation rate (marginal *R*^2^ = 0.30; conditional *R*^2^ = 0.03) for the most represented families of bees showed that the family Apidae was the main driver of the observed patterns. The contrasting differences between presence-absence and visitation rate for bees on PC1 (Fig. 3a and Fig. 3d) were driven by the family Andrenidae, which interacted more frequently on presence-absence interactions with plant species with low number of large flowers (Supplementary Fig. S8).

### Plant species functional roles

The variance of the different plant species-level network metrics was poorly explained by the three main axes of trait variation (Supplementary Fig. S9; interaction frequency ∼ PCs, conditional *R*^2^ = 0.11, marginal *R*^2^ = 0.02; normalized degree ∼ PCs, conditional *R*^2^ = 0.24, marginal *R*^2^ = 0.02; and, specialization ∼ PCs, conditional *R*^2^ = 0.37, marginal*R*^2^ = 0.03). Overall, the most notable trends were found on PC1 and PC3 for interaction frequency and specialization. On the flower number - flower size trade-off (PC1), interaction frequency was higher for plant species with more flowers but was lower for plant species with larger flowers (Supplementary Fig. S9a). On PC1, specialization showed the opposite trend (Supplementary Fig. S9g). On the style length trade-off (PC3), interaction frequency was lower for plants with shorter styles and lower autonomous selfing and higher for species with longer styles and higher autonomous selfing (Supplementary Fig. S9c). Again, specialization showed the opposite trend to interaction frequency (Supplementary Fig. S9i).

When we further investigated the combination of traits that drive plant network roles, we found that the regression tree for visitation frequency was best explained by plant height, nectar concentration and style length (Fig. 4a). Specifically, species taller than 3.9m had the highest interaction frequency, while species that were shorter than 3.9m and had a nectar concentration lower than 16% had the lowest interaction frequency. Normalized degree was best explained by nectar concentration, pollen grains per flower, plant height, flower width and autonomous selfing (Fig. 4b). Species with a nectar concentration over 49% had the highest levels of normalized degree, whereas species with nectar concentration lower than 49%, more than 21,000 pollen grains per flower and height less than 0.78m had the lowest normalized degree. Finally, specialization was best explained by plant height, ovule number, pollen grains per flower and autonomous selfing (Fig. 4c). Overall, plant species with the highest specialization were shorter than 1.3m, had more than 14,000 pollen grains per flower and autonomously self-pollinated less than 11% of their fruits. In contrast, species taller or equal than 5.1m and with lower than 14 ovules per flower had the lowest specialization values.

**Fig. 4.**
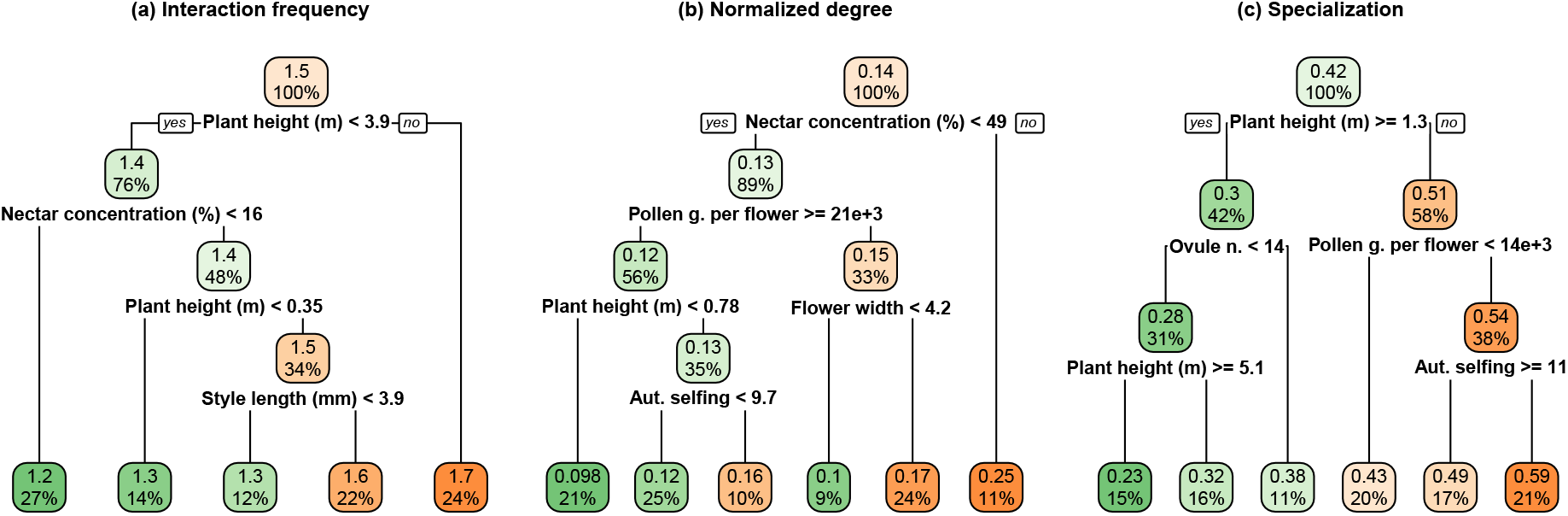
Contribution of traits in plant’s network roles. Regression tree analysis of interaction frequency (log-transformed), normalized degree and specialization for the subset of species with quantitative data for pollen and nectar traits. The superior value inside the node indicates the mean value of the different species-level metric and the lower value, the percentage of species that are considered in each node. Thus, the top node has the mean value of the named trait for the 100% of species. Each node has a yes/no question and when the condition is fulfilled, the branch turns to the ‘yes’ direction and when not, to the ‘no’ direction. This rationale is followed in all the regression trees as indicated in the first branch division of the topmost node of each tree.

## DISCUSSION

This study demonstrates that plant species exhibit clear trade-offs among their vegetative and reproductive traits and that these trade-offs determine interactions with floral visitors. These trade-offs are differentiated along three axes of trait variation: (i) flower number - flower size, (ii) pollinator dependence and (iii) style length. These reproductive trade-offs helped partly explain the presence of floral visitor interactions, but not their visitation rates. However, floral visitor guilds formed distinct relationships with the main axes of trait variation. Moreover, we found that the plant species functional roles within pollination networks were best explained by plant size and floral reward related traits.

Over half of all plant trait variation was captured by the flower number - flower size and pollinator dependence trade-offs. Trait variation on these two axes was associated with the ‘fast-slow continuum’ in plant^12^ and animal^45^ life-history strategies, as indicated by the different floral and reproductive biology traits associated with plant height, life form and lifespan. The ‘slow’ part of this continuum (i.e., tall trees and shrubs) included plant species with many flowers, few ovules, higher pollinator dependence, frequent occurrence of self-incompatibility and more complex breeding systems (e.g., monoecious and dioecious species). In contrast, plant species that employed the ‘fast’ strategy (i.e., short herbs), had fewer flowers, more ovules, frequent occurrence of self-compatibility and lower pollinator dependence. Further, on the first two axes of trait variation, we found additional support for the previously described positive association between higher outcrossing rate and larger floral display^32^. The positive correlation between larger floral display and higher pollinator dependence in our dataset further confirmed this trend (see Supplementary Fig. S10).

Despite the low predictive power of the main trait variation axes for broad-level interaction patterns (presence-absence of interactions and visitation rate), we found changes in the interaction patterns among and within floral visitor guilds across these axes that suggest plant life-history strategies influence plant-pollinator interactions. For example, all floral visitor guilds visited plant species with higher pollinator dependence more frequently, and high pollinator dependence was associated with large floral displays and greater pollen quantities (Fig. 1 and Supplementary Fig. S6). This trend is consistent with previous studies that show plant species with higher reproductive investment tend to be visited by pollinators more frequently^38,46,47^. In regard to the flower number - flower size and style length trade-offs, different pollinator guilds showed contrasting visitation rates across the continuum of trait variation, which could be associated with different pollination syndromes at a macroecological scale. For instance, bees and syrphid flies were clearly associated with opposing life-strategies on PC1 and PC3 (Fig. 3) suggesting possible niche partitioning^48,49^ between these two guilds. However, despite floral rewards not being included in the main analysis because there was insufficient data available, floral reward related traits were among the best at characterising species functional roles (Fig. 4). More detailed exploration of reproductive trade-offs in conjunction with floral rewards is needed to help elucidate plant-pollinator associations. In any case, it is worth noting that other local factors such as species relative abundances, surely explain part of the observed variability^17,50,51^ that reproductive trade-offs do not.

To conclude, we provide the first description of plant reproductive trade-offs using a large global dataset of plant traits. We identified the major reproductive strategies of flowering plants and how these strategies influence interactions with different floral visitor guilds. Although the explained variation that we found in the first two axes is lower than previous studies of vegetative traits^24,26^ it is consistent with the largest and most recent study that has characterised plant life strategies with vegetative and reproductive traits^12^. Future work needs to integrate the reproductive compromises that we have identified with vegetative and physiological trade-offs to create a more comprehensive spectrum of plant trait variation. Further, the varying level of phylogenetic signal among traits deserves further attention to understand evolutionary changes on mating and flower morphology in response to pollinators^52,53^. Finally, including plant-pollinator networks from unrepresented areas of the world and a more complete description of plant reproductive trade-offs is essential for a better understanding of the global patterns in plant-pollinator interactions.

## MATERIALS AND METHODS

### Plant-pollinator network studies

We selected 28 studies from 18 different countries that constituted a total of 64 plant-pollinator networks. These studies recorded plantpollinator interactions in natural systems and were selected so that we had broad geographical representation. Although these studies differ in sampling effort and methodology, all studies provided information about plant-pollinator interactions (weighted and non-weighted), which we used to build a database of plant species that are likely to be animal pollinated. Many of these networks are freely available either as published studies^54–56^ or available in online archives (e.g., The Web of Life^55^ and Mangal^57^). In total, our network dataset (see Supplementary Table S1) constituted 60 weighted (interaction frequency) and 4 unweighted (presence/absence of the interaction) networks, each sampled at a unique location and year, as well as eight meta-webs where interactions were pooled across several locations and multiple years.

### Taxonomy of plants and pollinators

All species names, genera, families and orders were retrieved and standardized from the taxonomy data sources NCBI (https://www.ncbi.nlm.nih.gov/taxonomy) for plants and ITIS (https://www.itis.gov/) for pollinators, using the R package *taxize*^58^ version *0*.*9*.*99*. We filled the ‘not found’ searches manually using http://www.theplantlist.org/ and http://www.mobot.org/ for plants and http://www.catalogueoflife.org/ for floral visitors.

### Functional traits

We selected 20 different functional traits based on their relevance to plant reproduction and data availability (Table 1). These included twelve quantitative and eight categorical traits belonging to three broader trait groupings (13 floral, 4 reproductive biology and 3 vegetative, Supplementary Information). For each plant species, we undertook an extensive literature and online search across a wide range of resources (plant databases, online floras, books, journals and images). From a total of 30,120 cells (20 columns × 1,506 species) we were able to fill 24,341 cells (80.8% of the dataset, see Supplementary Fig. S1 for missing values information for each trait).

**Table 1.** Quantitative and categorical traits used in this study.

| Quantitative traits |  | Categorical traits |  |  |
| --- | --- | --- | --- | --- |
| Type | Traits | Type | Traits | Categories |
| Vegetative | Plant height (m) | Vegetative | Lifepan | Short-lived<br>Perennial |
| Floral | Flower width (mm) | Vegetative | Life form | Herb<br>Shrub<br>Tree |
| Floral | Flower length (mm) | Floral | Flower shape | Brush<br>Campanulate<br>Capitulum<br>Open<br>Papilionaceous<br>Tube |
| Floral | Inflorescence width (mm) | Floral | Flower symmetry | Actinomorphic<br>Zygomorphic |
| Floral | Style length (mm) | Floral | Nectar | Presence<br>Absence |
| Floral | Ovules per flower | Reproductive biology | Autonomous selfing | None<br>Low<br>Medium<br>High |
| Floral | Flowers per plant | Reproductive biology | Compatibility system | Self-incomp.<br>Part. self-comp.<br>Self-comp. |
| Floral | Nectar (μl) | Reproductive biology | Breeding system | Hermaphrodite<br>Monoecious<br>Dioecious |
| Floral | Nectar (mg) |  |  |  |
| Floral | Nectar concentration (%) |  |  |  |
| Floral | Pollen grains per flower |  |  |  |
| Reproductive biology | Autonomous selfing (fruit set) |  |  |  |

### Phylogenetic Distance

We calculated the phylogenetic distance between different plant species using the function *get_tree* from the package *rtrees* (https://github.com/daijiang/rtrees), which downloads phylogenetic distances from the extended R implementation of the Open Tree of Life^59,60^.

### Data Imputation

Trait missing values were imputed with the function *missForest*^61^ which allows imputation of data sets with continuous and categorical variables. We accounted for the phylogenetic distance among species on the imputation process by including the eigenvectors of a principal component analysis of the phylogenetic distance (PCoA) which has been shown to improve the performance of *missForest*^62^. To extract the eigenvectors, we used the function *PVRdecomp* from the package *PVR*^63^ based on a previous conceptual framework that considers phylogenetic eigenvectors^64^. Although the variable of autonomous selfing had a high percentage of missing values (68%), we were able to solve this by back transforming the qualitative column of autonomous selfing to numerical. The categories of ‘none’, ‘low’, ‘medium’ and ‘high’ were converted to representative percentages of each category 0%, 13%, 50.5% and 88% respectively. This reduced the percentage of missing values for this column from 68% to 35% and allowed the imputation of this variable. However, we were unable to include nectar and pollen traits on the imputation process because of the high percentage of missing values (Supplementary Fig. S1). Hence, the imputed dataset had 1,506 species, seven categorical and eight numerical variables and 5.79% of missing values. Further, we conducted an additional imputation process on the subset of species with data for pollen per flower and microliters of nectar. This subset comprised 755 species, 8.01% missing values and all traits but milligrams of nectar (∼50% of missing values) were included in the imputation process.

### Plant strategies

We explored the trade-offs between different quantitative plant functional traits with a phylogenetically informed Principal Component Analysis (pPCA). We did not include the quantitative variables of flower length and inflorescence width because they were highly and moderately correlated to flower width respectively (Pearson’s correlation = 0.72, *P* < 0.01 and Pearson’s correlation = 0.36, *P* < 0.01), and thus we avoided overemphasizing flower size on the spectrum of trait variation. Although qualitative traits were not included in the dimensionality reduction analysis, we also investigated the association of the different qualitative traits with the main axes of trait variation. Prior to the analyses, we excluded outliers and standardized the data. Due to the high sensitivity of dimensionality reduction to outliers, we excluded values within the 2.5th–97.5th percentile range^65^, and thus our final dataset had 1,236 species. Then, we log transformed the variables to reduce the influence of outliers and z-transformed (X= 0, SD=1) so that all variables were within the same numerical range. We performed the pPCA using the function *phyl*.*pca* from the package *phytools*^66^ (version *0*.*7-70*) with the method lambda (*λ*) that calculates the phylogenetic correlation between 0 (phylogenetic independence) and 1 (shared evolutionary history) and we implemented the mode covariance because values for each variables were on the same scale following transformation^67^. Moreover, to corroborate that our imputation of missing values did not affect our results, we conducted a pPCA on the full dataset without missing values (see Supplementary Fig. S2). We found little difference between the explained variance with the imputed dataset (51.08%) and the dataset without missing values (52.87%). In addition, the loadings on each principal component had a similar contribution and correlation patterns, with the exception of plant height which showed slight variations between the imputed and non-imputed dataset. Finally, we conducted an additional phylogenetic informed principal component analysis for the subset of species with pollen and nectar quantity. For this, we included all quantitative traits considered in the main pPCA plus pollen grains and microlitres of nectar per flower.

### Phylogenetic signal of traits

We calculated the phylogenetic signal of the different quantitative traits on the imputed dataset with the full set of species (N = 1,506) with the package *phytools*^66^ version *0*.*7-70* and we used Pagel’s *λ* as a measurement of the phylogenetic signal. However, for pollen and nectar traits, phylogenetic signal was calculated only on the subset of species that had quantitative information for these traits (N = 755).

### Network analyses

Analyses were conducted on the subset of 60 weighted networks sampled in a unique flowering season and site, which included 556 plant and 1,126 pollinator species. These networks were analysed in their qualitative (presence-absence) and quantitative (interaction frequency) form. First, we analysed the binary version of these weighted networks with presence-absence information that assumes equal weight across interactions. Second, we analysed the untransformed weighted networks with interaction frequency that accounts for the intensity of the interaction. Although floral visitors are not always pollinators and interaction frequency does not consider each pollinator species efficiency^68^, interaction frequency can provide valuable information of the contribution of floral visitors to pollination^69,70^. In total, our network dataset (excluding meta-webs and non-weighted networks) included 2,256 interactions of bees with plants, 1,768 non-syrphid-Diptera interactions, 845 syrphids interactions, 437 Lepidoptera interactions, 432 Coleoptera interactions and 362 non-bee-Hymenoptera interactions. Sampling methods varied across networks but this was accounted for in analyses by considering them in the random effects of the modelling process. All analyses were conducted in R version *4*.*0*.*3*.

### Visitation patterns

We used Bayesian modelling (see below for details) to explore the effect of floral visitor groups and the main axes of trait variation (pPCA with imputed dataset) on both qualitative (presence/absence) and quantitative (visitation rate) floral interactions per plant species. For this, we divided floral visitors into six main guilds that differ in life form, behaviour and are likely to play a similar ecological role: (i) bees (Hymenoptera-Anthophila), (ii) non-bee-Hymenoptera (Hymenopteranon-Anthophila), (iii) syrphids (Diptera-Syrphidae), (iv) non-syrphid-Diptera (Dipteranon-Syrphidae), (v) Lepidoptera and (vi) Coleoptera. Moreover, because the guild of bees was the most represented group with 2,256 records and had the highest frequency of visits of all groups, we also explored the presence-absence of interaction and visitation rate of the main bee families (Andrenidae, Apidae, Colletidae, Halictidae and Megachilidae) on the trait space. In addition, we found that *Apis mellifera* was the floral visitor with the largest proportion of records counted (7.55% of the total). This finding is consistent with previous research showing that *A. mellifera* was the most frequent floral visitor in a similar dataset of 80 plant-pollinator networks in natural ecosystems^71^. Hence, to control for the effect of *A. mellifera* on the observed visitation patterns of bees, we conducted an analogous analysis with presence-absence of interaction and visitation rate excluding *A. mellifera*. We found that *A. mellifera*, was partly driving some of the observed trends on PC1 (Supplementary Fig. S3). However, we did not detect major differences on PC2 and PC3.

We implemented Bayesian generalized linear mixed models using the R package *brms*^72^ (version *2*.*14*.*6*). We modelled the frequency of visits as a function of the main axes of plant trait variation and their interactions with floral visitor functional groups (Visits ∼ PC1 x FGs + PC2 x FGs + PC3 x FGs). Because we were interested in possible differences in the visitation patterns among floral visitors groups to plants with different strategies, we included interactions between the main axes of trait variation (PC1, PC2 and PC3) and the floral visitor guilds. In this model, we added a nested random effect of networks nested within the study system to capture the variation in networks among studies and within networks. Moreover, we included the phylogenetic covariance matrix as a random factor due to the possible shared evolutionary histories of species and therefore lack of independence across them. We specified this model with a zero inflated negative binomial distribution and weakly informative priors from the brms function. We run this model for 3,000 iterations and with previous 1,000 warm up iterations. We set delta (Δ) to 0.99 to avoid divergent transitions and visualized the posterior predictive checks with the function *pp_check* using the *bayesplot* package^73^ (version *1*.*7*.*2*).

### Plant species functional roles

We investigated whether different quantitative traits determined plant species functional roles using Bayesian modelling and regression trees. For this, we selected simple and complementary species-level network metrics commonly applied in bipartite network studies^74^ with a straightforward ecological interpretation relevant to our research goals. The different plant species-level metrics were: (i) sum of visits per plant species; (ii) normalized degree, calculated as the number of links per plant species divided by the total possible number of partners; and (iii) specialization (d’)^75^, which measures the deviation of an expected random choice of the available interaction partners and ranges between 0 (maximum generalization) and 1 (maximum specialization). Normalized degree and specialization were calculated with the *specieslevel* function from the R package *bipartite*^74^ (version *2*.*15*).

First, we modelled the distinct plant species metrics (sum of visits, normalized degree and plant specialization) as a function of the three main axes of trait variation (plant species level metric ∼ PC1 + PC2 + PC3). For each response variable (i.e., each plant species level metric), we used different distribution families (zero inflated negative binomial for the sum of visits, weibull for normalized degree and zero one inflated beta for specialization). Finally, we used the same random factors, model settings and conducted the same posterior predictive checks for each model as detailed above in the ‘visitation patterns section’.

Second, to better understand these complex trait relationships, we used regression trees. Regression trees are recursive algorithms which can detect complex relationships among predictors and allow identification of the relevance of specific trait combinations on species functional roles. We focused exclusively on quantitative traits because almost all categorical traits were statistically associated with the first two axes of trait variation (Supplementary Table S2). We conducted this analysis using the *rpart* package^76^ version *4*.*1-15* with method *‘anova’* with a minimum of 50 observations per terminal node and we used the *rpart*.*plot* package^77^ version *3*.*0*.*9* to plot the regression trees. We considered the species level indices as response variables (interaction frequency, normalized degree and specialization) and we performed one regression tree per metric using the different quantitative traits as predictors. We calculated two regression trees per plant specieslevel metric, one for the full set of species and another for the subset of species for which we had pollen and nectar traits. We focused on regression trees that included floral rewards because they consistently showed pollen and nectar traits as being the best for explaining the different species-level metrics (see Supplementary Fig. S4).

## Acknowledgements

This is study was supported by the project SAFEGUARD (101003476 H2020-SFS-2019-2). We thank all researchers that made their data available for our analysis. We thank Bryony Wilcox, Greg Bible, Mercedes Sanchez-Lanuza and David Ragel for their help with data collection. We also thank Jason Tylianakis for his comments on the manuscript before submission. JBL thanks the University of New England for the funding provided to carry out this work.

