## Supplementary information for "Trade-offs among plant reproductive traits determine interactions with floral visitors"

### Contents:

##### Supplementary text

Description of the traits compiled in this study with an illustrative figure of the main flower morphologies (Supplementary text Fig 1)

##### Supplementary Tables

Supplementary Table S1: List of the 28 plant-pollinator studies used to build the plant trait database

Supplementary Table S2: Loadings of the first three axes of trait variation of the phylogenetic informed principal component analysis with the full set of species

Supplementary Table S3: Loadings of the first three axes of trait variation of the phylogenetic informed principal component analysis with the subset of species with data of nectar and pollen quantity

Supplementary Table S4: Statistical association between the different categorical variables and the fisrt three main axes of trait variation with the full set of species

Supplementary Table S5: Phylogenetic signal of the different quantitative traits

##### Supplementary Figures

Supplementary Fig. S1: Percentage of present and missing values of the different plant traits

Supplementary Fig. S2: Phylogenetic informed principal component analysis for the species that did not have missing values

Supplementary Fig. S3: Fitted posterior estimates of the number of visits made by bees including and exluding honey bees on the main axes of trait variation

Supplementary Fig. S4: Regression tree analyses

Supplementary Fig. S5: PC1 and PC3 of the phylogenetically informed principal component analysis with the full set of species

Supplementary Fig. S6: Phylogenetic informed principal component analysis for the subset of species with floral rewards.

Supplementary Fig. S7: Statistical comparison of the different categories of qualitative traits on the main two axes of trait variation with the full set of species

Supplementary Fig. S8: Presence-absence of interaction and visitation rate of the main bee families on the main axes of trait variation

Supplementary Fig. S9: Plant species level metrics across the three main axes of trait variation

Supplementary Fig. S10: Pearson correlation between the product of flower number-flower size and autonomous selfing level

#### Description of the traits compiled in this study

*Reproductive traits*

- Breeding system: The different plant species were classified in hermaphrodite, dioecious and monoecious species. Intermediate breeding systems or more complex ones were also annotated but all the species were divided into these three main categories for simplicity of the analysis.
- Selfing level: We recorded the selfing level of the different species with both quantitative and qualitative data. The qualitative data was divided in four main categories, high selfers, medium selfers, low selfers and none which was for the species that were unable to self-pollinate. In addition, quantitative data was also divided into these four categories with the following criteria: from 0% to lower than 1% 'none', from 1% to 25% 'low', from 26% to 75% 'medium' and from 76% to 100% 'high'.
- Compatibility system: The different species were divided in three main categories in order to know their ability to self-pollinate. These categories were self-compatible, partially self-compatible and self-incompatible species. The field of selfing level is partly complementary to the compatibility system but is important to note that not all the self-compatible species or partially self-compatible species will self-pollinate.

*Floral traits*

- Flower symmetry: We also recorded if the flowers were bilaterally symmetrical "zygomorphic" or were radially symmetrical "actinomorphic".
- Flower and inflorescence size: We searched for flower length and flower and inflorescence width (mm) for all species. When possible, we calculated the not found measurements with the help of online images and the software ImageJ.
- Flower number per plant: We compiled information about the number of flowers per plant for all species. However, this field was rare to find and we also used online images of the different species in order to calculate rough numbers of flowers per plant. We referenced all the filled fields in order to be able to follow the images that were used for these counts. It is important to note, that these numbers are not pretended to be the exact number of flowers per species but an approximate indicator of the reproductive investment for the different species that allow the macroecological analysis of this field.
- Ovule number: We searched for the number of ovules per flower for all the different species. The number of ovules of Asteraceae species are considered as the total number of ovules per capitulum. Many species were filled by genus or family level because the number of ovules in this taxonomic groups is considered to be mostly constant (e.g., Lamiaceae Boraginaceae or Apiaceae).
- Flower morphology: We looked for images and illustrations of the flowers from the different species on the floras and available resources in order to categorize the flower shape. We divided the flowers in 8 main different categories: open, bowl, tube, campanulate, funnelform, papilionaceous, brush and capitulum. However, we ended

up groupping all flowers on the following 6 morphological groups for analyses: tube, papilonaceous, open, capitulum, campanulate and brush (Supplementary text Fig. 1).

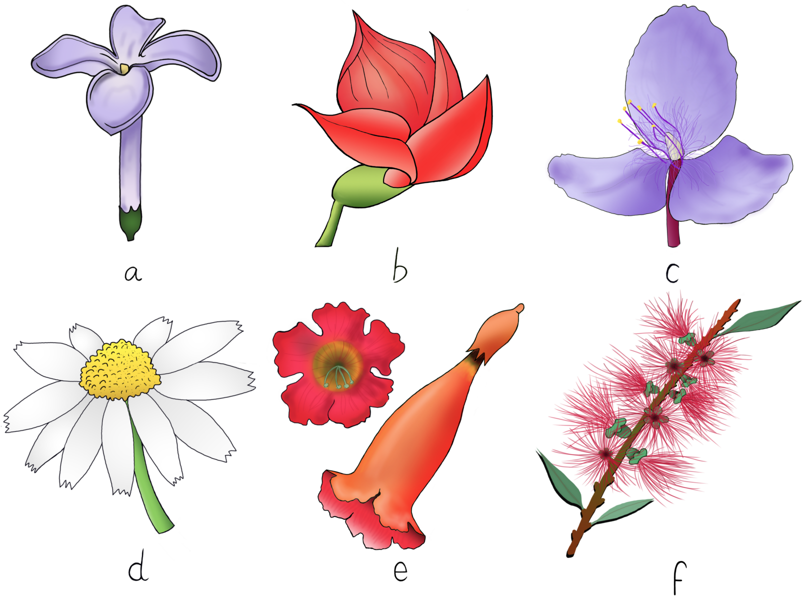


**Supplementary text Fig. 1 | Main floral morphologies considered in this study.** All species were groupped within these categories for analysis: (a) tube, (b) papilonaceous, (c) open, (d) capitulum, (e) campanulate and (f) brush.

- Style length: The length of the style (mm) of the different species was also included. When possible, we calculated style length from images and illustrations on the online floras.
- Nectar provision: We recorded the presence and absence of nectar for all species. In addition, we also searched for microlitres and milligrams of nectar per flower and nectar concentration. In general terms, nectar data was rarely present and we tried to minimise the loss of information by filling some species at family level. For instance, Solanaceae species are described as nectarless, and species that belonged to this family were recorded with 'absence' of nectar.
- Pollen grains per flower: We recorded pollen grains per flower for all species but this field was rarely described in the literature. When just the pollen:ovule ratio was found we converted it back to pollen grains by multiplying it by the average ovule number found for each species.

*Vegetative traits*

- Life form, life span and plant height: We divided the different plant species in 4 main categories: herbs, vines, shrubs and trees. Moreover, we also divided the species between short-lived species (annual, biennial and short-lived perennials) and perennial species (long-lived). Finally, we also searched for the average height (m) of the different species and annotated the maximum and minimum height when possible. We conducted the average between the maximum and minimum height to get an approximate average height of the species when the average was not indicated.

**Supplementary Table S1 | List of the 28 plant-pollinator studies used to build the plant trait database.** Each study is shown with the first author that conducted the study, number of networks or metawebs that contains, type of information that contains (weighted or unweighted), the structure (web or metaweb), year of publication and digital object identifier or permanent link for each study.

| First author | Year | Web N. | Network type | DOI |
|---|---|---|---|---|
| Arroyo-Correa | 2019 | 3 | Weighted web | https://doi.org/10.1111/1365-2745.13332 |
| Bartomeus | 2008 | 6 | Weighted web | https://doi.org/10.1007/s00442-007-0946-1 |
| Bartomeus | 2008 | 16 | Weighted web | https://github.com/ibartomeus/BeeFunData |
| Bek | 2006 | 1 | Unweighted web | Unpublished, Master thesis |
| Bundgaard | 2003 | 1 | Weighted web | Unpublished, Master thesis |
| Burkle | 2013 | 1 | Weighted web | https://doi.org/10.1126/science.1232728 |
| Dicks | 2002 | 2 | Weighted web | https://doi.org/10.1046/j.0021-8790.2001.00572.x |
| Dupont | 2003 | 3 | Weighted web | https://doi.org/10.1111/j.1365-2656.2008.01501.x |
| Elberling | 1999 | 1 | Weighted web | https://doi.org/10.1111/j.1600-0587.1999.tb00507.x |
| Fang | 2008 | 1 | Weighted web | https://doi.org/10.1111/1749-4877.12190 |
| Inouye | 1988 | 1 | Weighted web | https://doi.org/10.1111/j.1442-9993.1988.tb00968.x |
| Inouye | 1990 | 1 | Weighted metaweb | http://hdl.handle.net/2433/156099 |
| Kaiser-Bunbury | 2017 | 8 | Weighted web | https://doi.org/10.1038/nature21071 |
| Kaiser-Bunbury | 2011 | 6 | Weighted web | https://doi.org/10.1111/j.1365-2745.2010.01732.x |
| Kaiser-Bunbury | 2010 | 2 | Weighted web | https://doi.org/10.1016/j.ppees.2009.04.001 |
| Kato | 2000 | 1 | Unweighted web | http://hdl.handle.net/2433/156116 |
| Kevan | 1970 | 1 | Unweighted web | https://doi.org/10.2307/2258569 |
| Lundgren | 2005 | 1 | Weighted web | https://doi.org/10.1657/1523-0430(2005)037[0514:TDAHCW]2.0.CO;2 |

*(continued)*

| First author | Year | Web N. | Network type | DOI |
|---|---|---|---|---|
| McMullen | 1993 | 1 | Unweighted metaweb | https://biostor.org/reference/244737 |
| Olesen | 2002 | 2 | Weighted web | https://doi.org/10.1046/j.1472-4642.2002.00148.x |
| Peralta | 2006 | 4 | Weighted web | https://doi.org/10.1111/ele.13510 |
| Primack | 1983 | 3 | Unweighted metaweb | https://doi.org/10.1080/0028825X.1983.10428561 |
| Ramirez | 1989 | 1 | Unweighted web | https://doi.org/10.2307/2388282 |
| Ramirez | 1992 | 1 | Unweighted metaweb | https://doi.org/10.1111/j.1095-8339.1992.tb00294.x |
| Robertson | 1929 | 1 | Unweighted metaweb | https://doi.org/10.5962/bhl.title.11538 |
| Small | 1976 | 1 | Weighted web | /13960/t4km08d21 |
| Souza | 2017 | 1 | Weighted web | https://doi.org/10.1111/1365-2745.12978 |
| Traveset | 2013 | 1 | Weighted metaweb | https://doi.org/10.1098/rspb.2012.3040 |

**Supplementary Table S2 | Loadings of the first three axes of trait variation of the phylogenetic informed principal component analysis with the full set of species.**

| | PC1 | PC2 | PC3 |
|---|---|---|---|
| Autonomous selfing | 0.03 | 0.85 | -0.51 |
| Flowers per plant | 0.75 | -0.27 | -0.24 |
| Flower width | -0.67 | -0.38 | -0.30 |
| Style length | -0.34 | -0.37 | -0.66 |
| Ovule number | -0.53 | 0.00 | -0.02 |
| Plant height | 0.56 | -0.40 | -0.46 |
| Explained variation | 26.72 | 25.08 | 19.17 |

**Supplementary Table S3 | Loadings of the first three axes of trait variation of the phylogenetic informed principal component analysis with the subset of species with data of nectar and pollen quantity.**

| | PC1 | PC2 | PC3 |
|---|---|---|---|
| Autonomous selfing | 0.19 | 0.76 | 0.35 |
| Flowers per plant | -0.62 | -0.35 | 0.45 |
| Flower width | 0.66 | -0.44 | -0.08 |
| Style length | 0.54 | -0.42 | 0.13 |
| Ovule number | 0.60 | 0.00 | -0.04 |
| Plant height | -0.38 | -0.52 | 0.53 |
| Microlitres of Nectar per flower | 0.51 | 0.06 | 0.66 |
| Pollen per flower | 0.16 | -0.50 | -0.11 |
| Explained variation | 23.40 | 21.67 | 14.36 |

**Supplementary Table S4 | Statistical association between the different categorical variables and the first three main axes of trait variation with the full set of species.**

| Functional traits | Sum Sq | F value | Pr(>F) | PC |
|---|---|---|---|---|
| Breeding system | 304.59 | 119.50 | 0.00 | PC1 |
| Compatibility system | 89.12 | 23.31 | 0.00 | PC1 |
| Lifespan | 35.65 | 27.97 | 0.00 | PC1 |
| Life form | 565.87 | 222.00 | 0.00 | PC1 |
| Flower shape | 132.24 | 20.75 | 0.00 | PC1 |
| Flower symmetry | 0.37 | 0.29 | 0.59 | PC1 |
| Nectar provision | 0.38 | 0.29 | 0.59 | PC1 |
| Breeding system | 304.59 | 119.50 | 0.00 | PC2 |
| Compatibility system | 89.12 | 23.31 | 0.00 | PC2 |
| Lifespan | 35.65 | 27.97 | 0.00 | PC2 |
| Life form | 565.87 | 222.00 | 0.00 | PC2 |
| Flower shape | 132.24 | 20.75 | 0.00 | PC2 |
| Flower symmetry | 0.37 | 0.29 | 0.59 | PC2 |
| Nectar provision | 0.38 | 0.29 | 0.59 | PC2 |

**Supplementary Table S5 | Phylogenetic signal of the different quantitative traits.**

| Functional traits | Lambda | P-value |
|---|---|---|
| Autonomous selfing | 0.34 | 0.00 |
| Flower number | 0.69 | 0.00 |
| Inflorescence width | 0.57 | 0.00 |
| Flower width | 0.73 | 0.00 |
| Flower length | 0.75 | 0.00 |
| Style length | 0.49 | 0.00 |
| Ovule number | 1.00 | 0.00 |
| Plant height | 0.96 | 0.00 |
| Nectar per flower ($\mu$l) | 0.14 | 0.00 |
| Nectar concentration (%) | 0.65 | 0.00 |
| Pollen grains per flower | 1.00 | 0.00 |

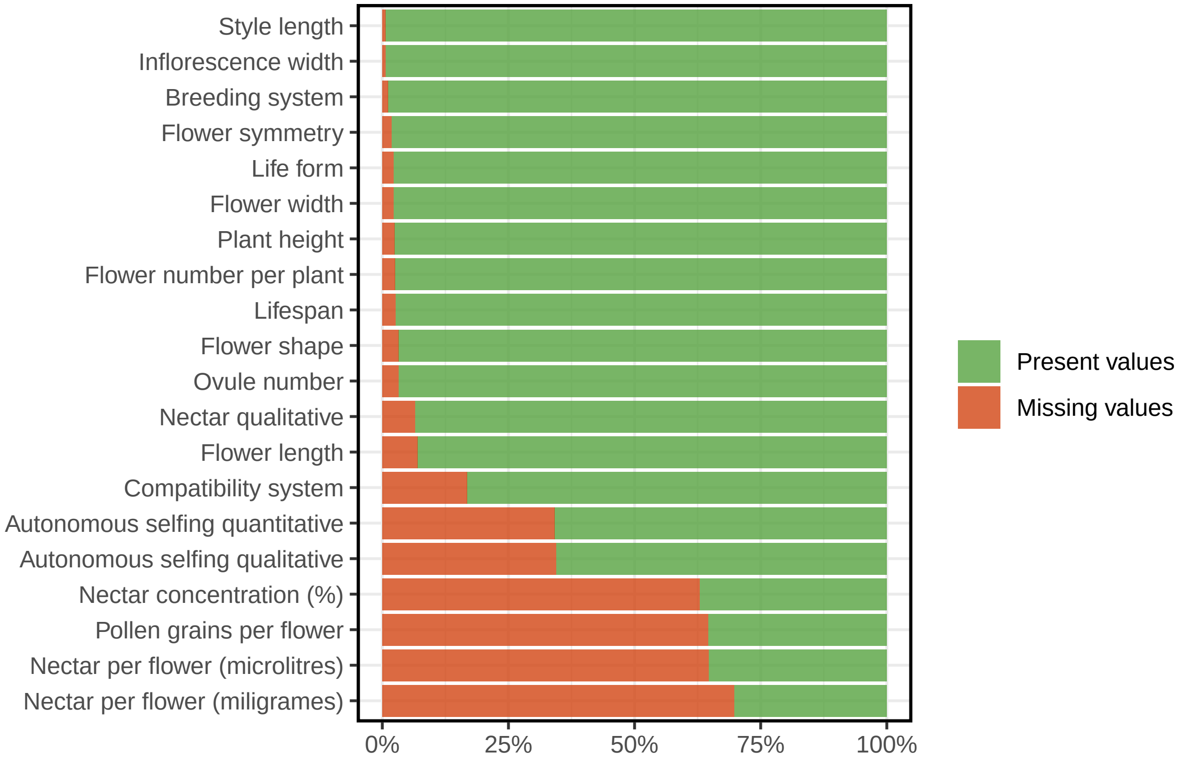


**Supplementary Fig. S1 | Percentage of present and missing values of the different plant traits.**

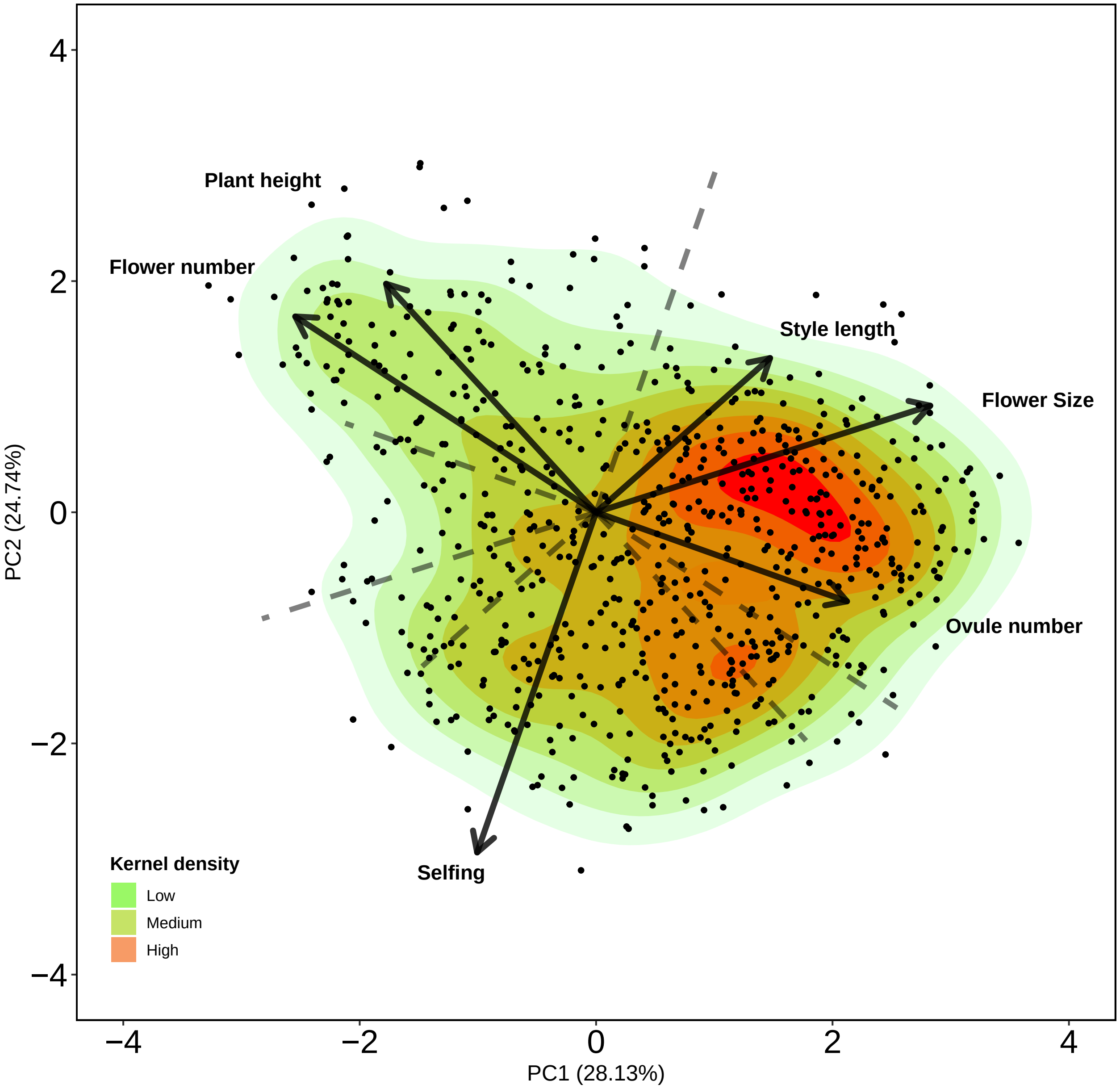


**Supplementary Fig. S2 | Phylogenetic informed principal component analysis for the species that did not have missing values.**

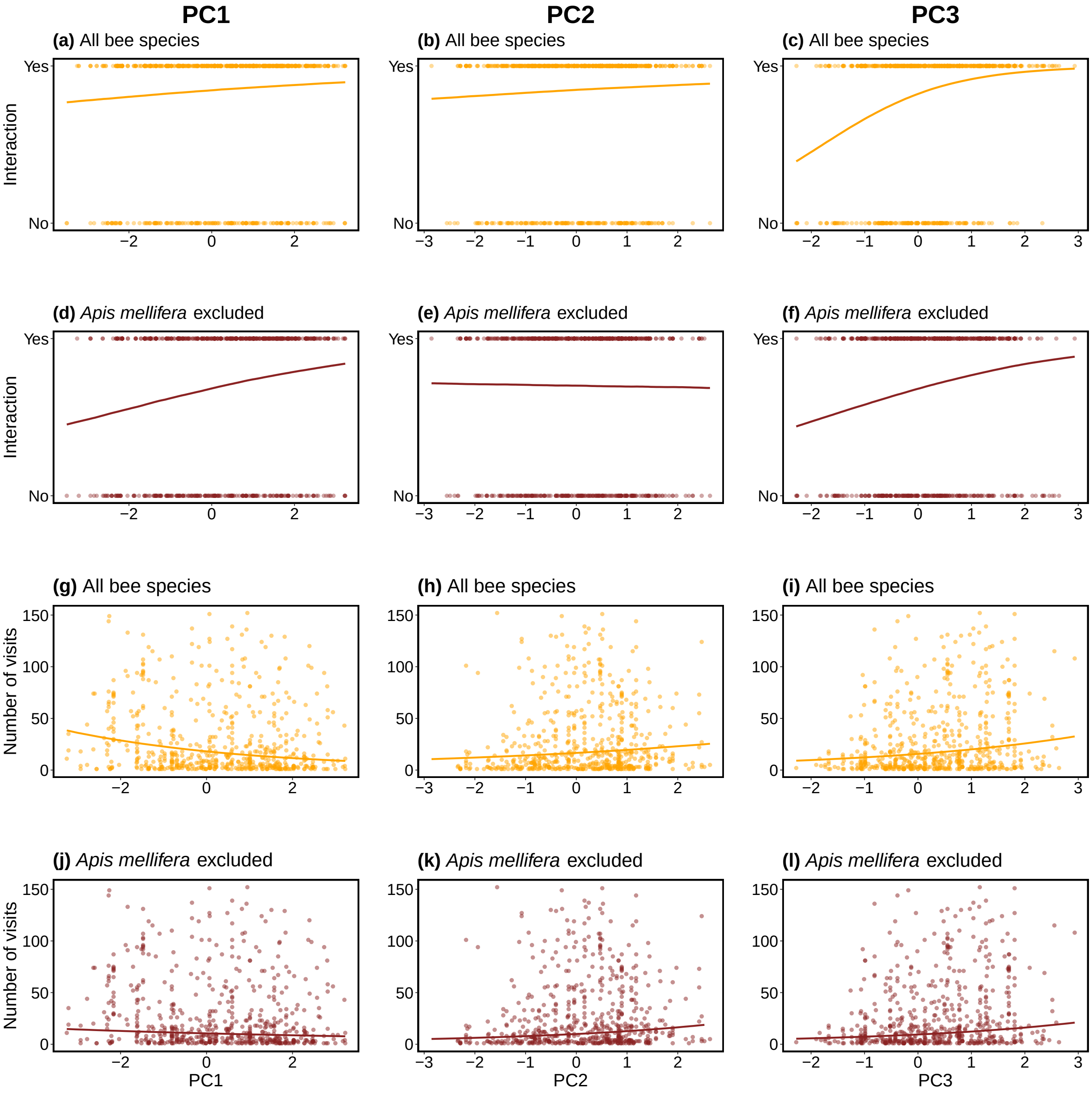


**Supplementary Fig. S3 | Fitted posterior estimates of the interaction (yes/no) and number of visits made by bees including and exluding honey bees on the main axes of trait variation.** The superior panel (plots a, b c, d, e and f) shows the comparison for presence-absence and the lower panel (plots g, h, i, j, k and l) the comparison for visitation rates.

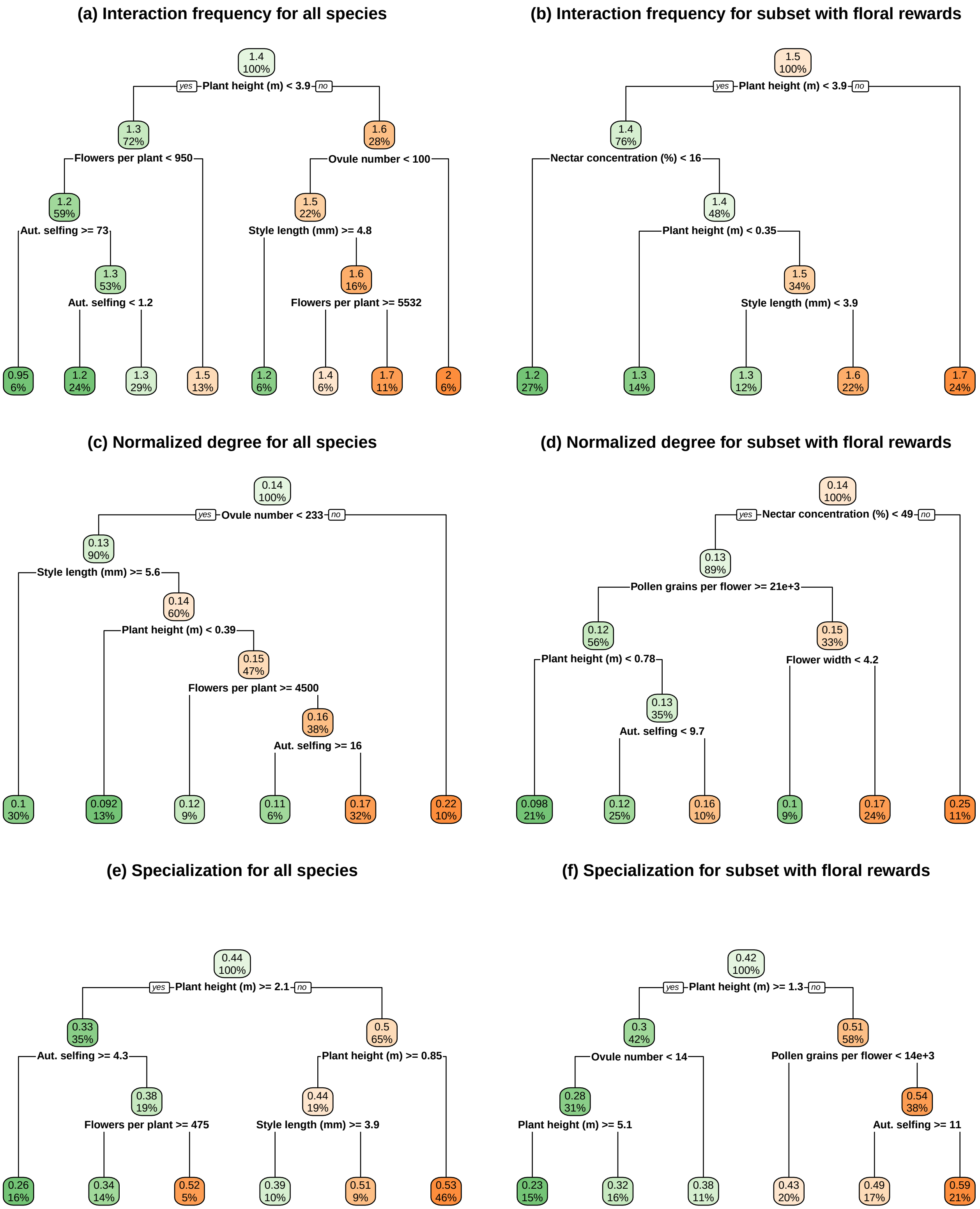


**Supplementary Fig. S4 | Regression tree analyses.** Comparison of the regression tree analyses between the full set of species and the subset of species with floral rewards for each plant species level metric.

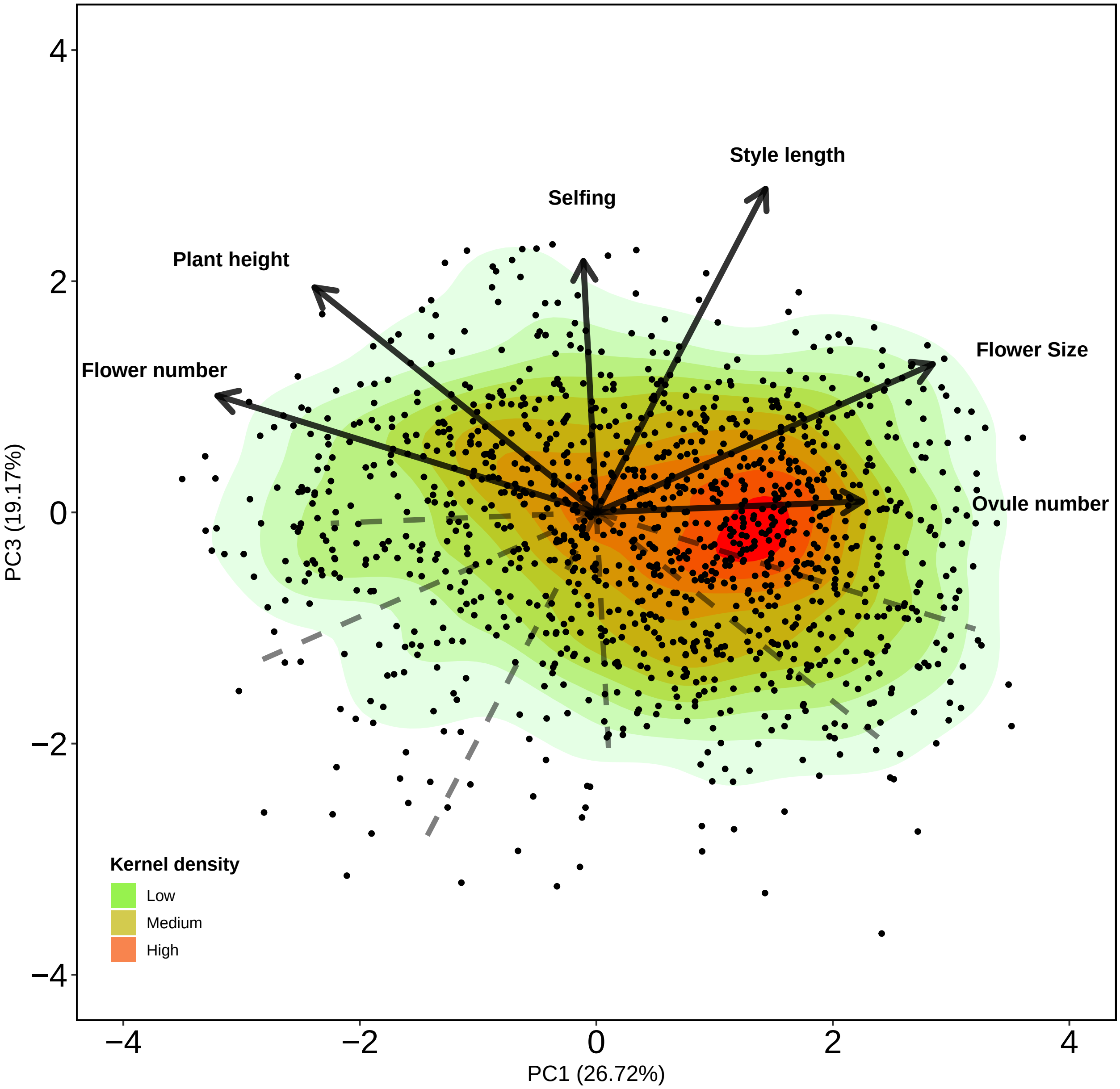


**Supplementary Fig. S5 | PC1 and PC3 of the phylogenetically informed principal component analysis with the full set of species.**

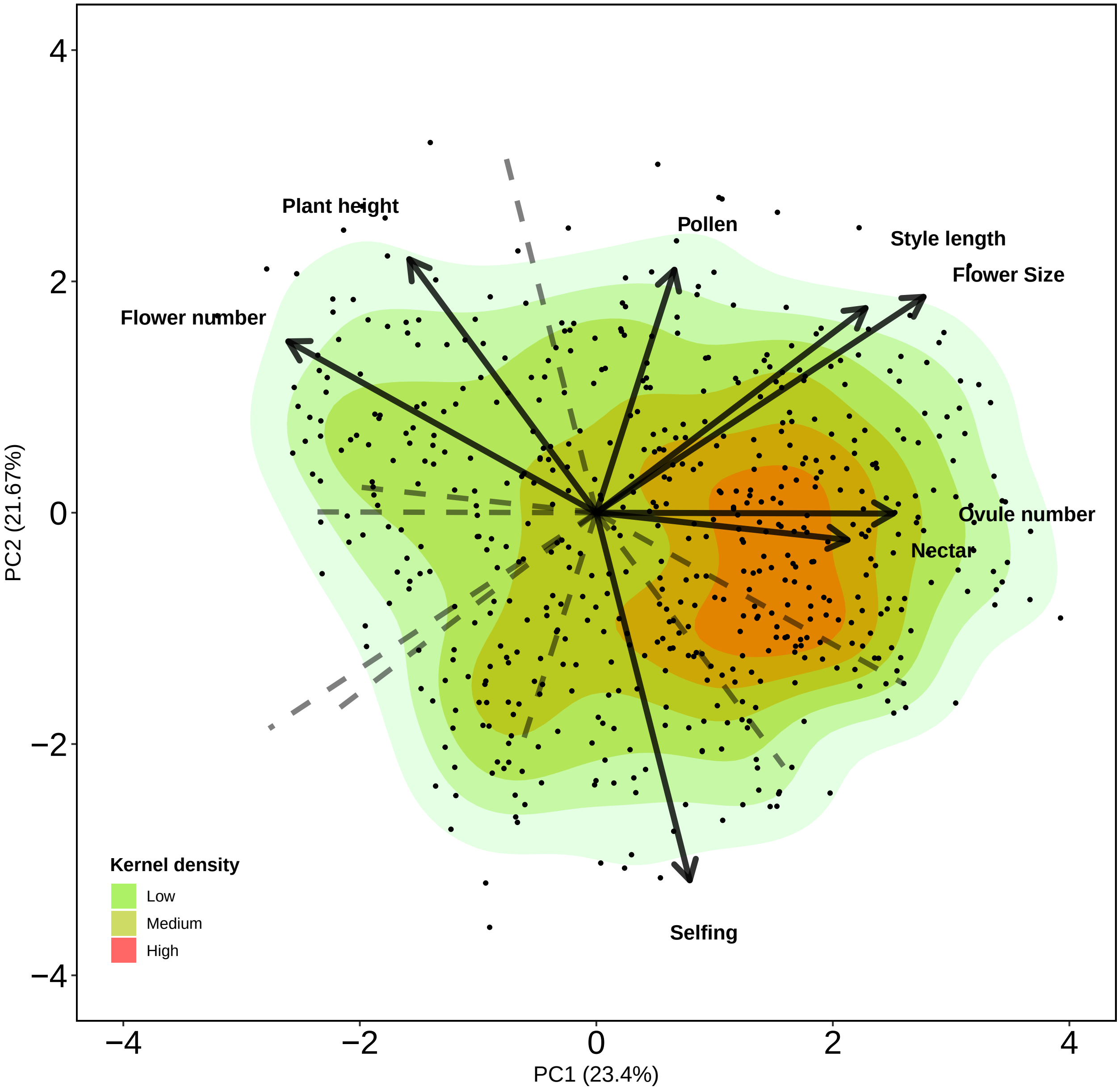


Supplementary Fig. S6 | Phylogenetic informed principal component analysis for the subset of species with floral rewards.

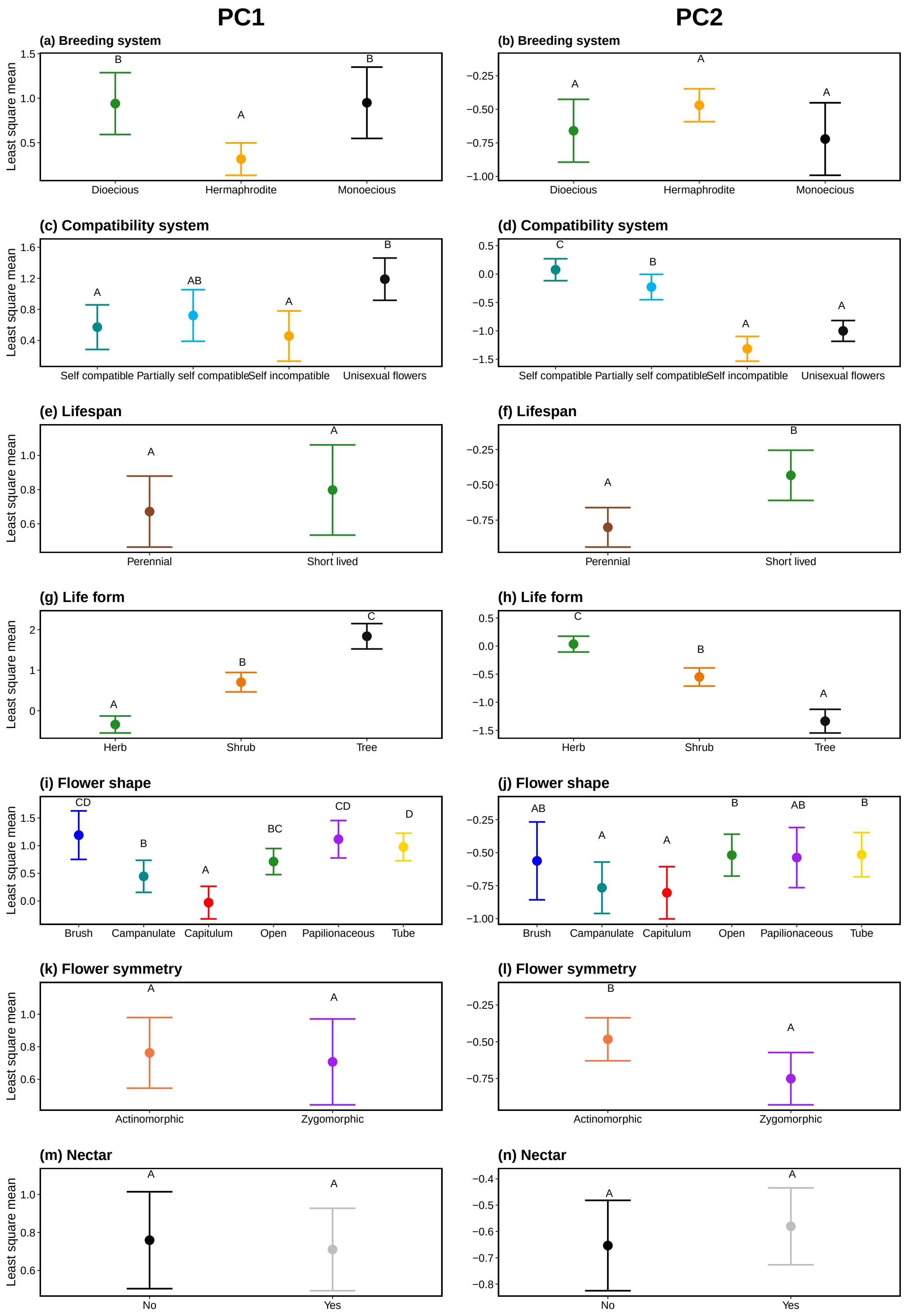


**Supplementary Fig. S7 | Statistical comparison of the different categories of qualitative traits on the main two axes of trait variation with the full set of species.** Categories that differ significantly are denoted with a different letter.

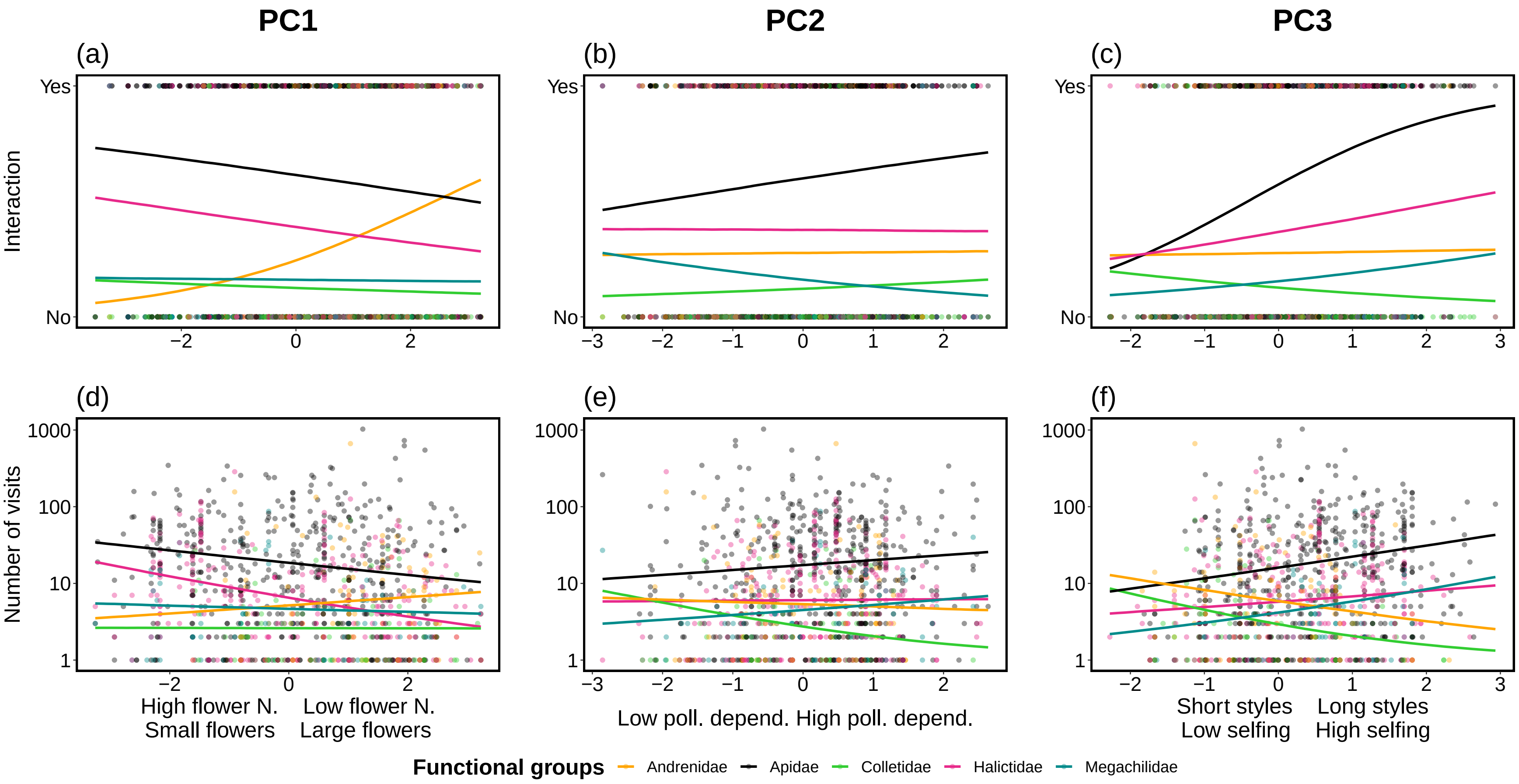


**Supplementary Fig. S8 | Interaction (yes/no) and visitation rate of the main bee families on the main axes of trait variation.** Fitted posterior estimates of the presence-absence of interaction (plots a, b and c) and visitation rate (plots e, f and g) of Andrenidae, Apidae, Colletidae, Halictidae and Megachilidae across PC1, PC2 and PC3. For visualization purposes, the response variable of number of visits was log-transformed (Y-axis of lower panel)

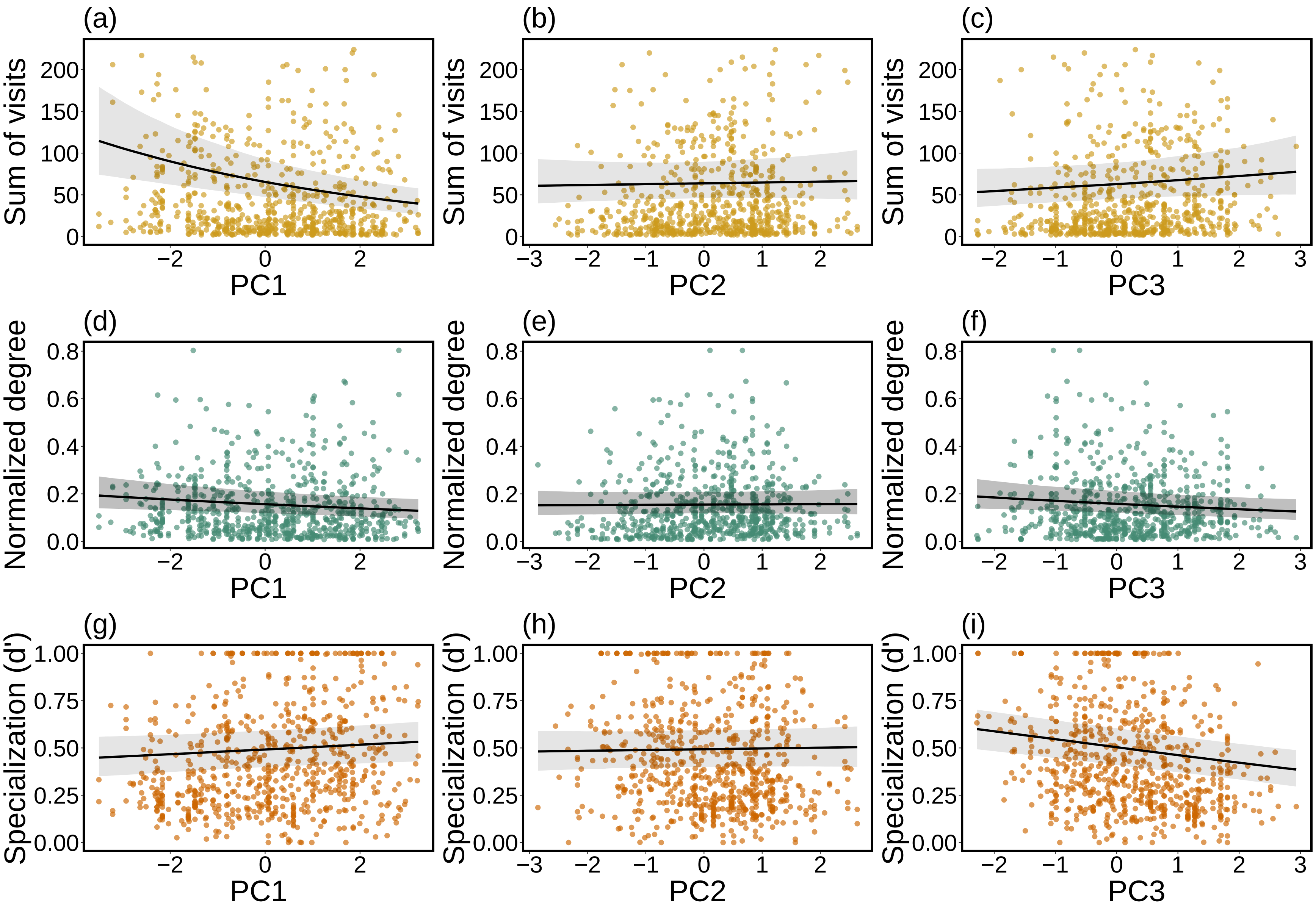


**Supplementary Fig. S9 | Plant species level metrics across the three main axes of trait variation.**

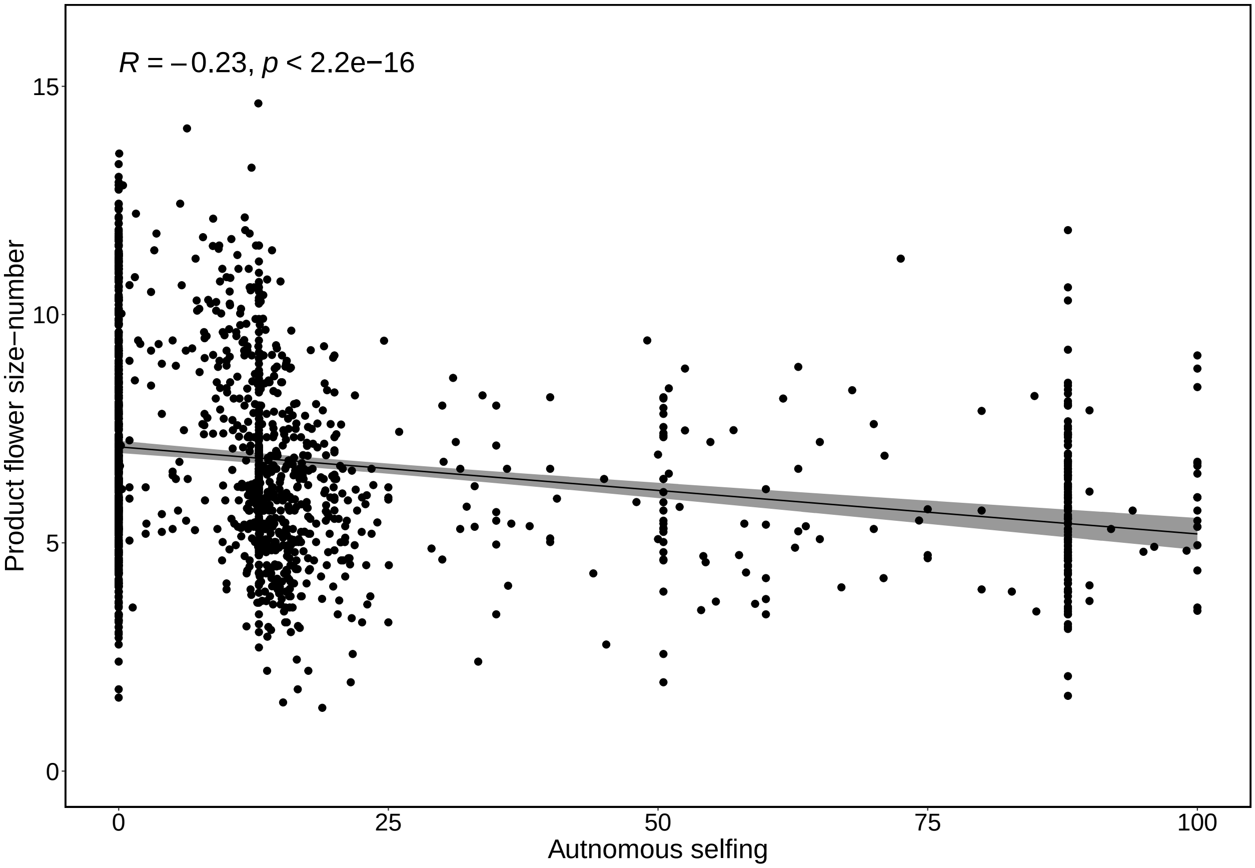


**Supplementary Fig. S10 | Pearson correlation between the product of flower number-flower size and autonomous selfing level.**
